# Automated machine learning and explainable AI (AutoML-XAI) for metabolomics: improving cancer diagnostics

**DOI:** 10.1101/2023.10.26.564244

**Authors:** Olatomiwa O. Bifarin, Facundo M. Fernández

## Abstract

**Motivation:** Metabolomics generates complex data necessitating advanced computational methods for generating biological insight. While machine learning (ML) is promising, the challenges of selecting the best algorithms and tuning hyperparameters, particularly for nonexperts, remain. Automated machine learning (AutoML) can streamline this process; however, the issue of interpretability could persist. This research introduces a unified pipeline that combines AutoML with explainable AI (XAI) techniques to optimize metabolomics analysis.

**Results:** We tested our approach on two datasets: renal cell carcinoma (RCC) urine metabolomics and ovarian cancer (OC) serum metabolomics. AutoML, using auto-sklearn, surpassed standalone ML algorithms such as SVM and random forest in differentiating between RCC and healthy controls, as well as OC patients and those with other gynecological cancers (Non-OC). Autosklearn employed a mix of algorithms and ensemble techniques, yielding a superior performance (AUC of 0.97 for RCC and 0.85 for OC). Shapley Additive Explanations (SHAP) provided a global ranking of feature importance, identifying dibutylamine and ganglioside GM(d34:1) as the top discriminative metabolites for RCC and OC, respectively. Waterfall plots offered local explanations by illustrating the influence of each metabolite on individual predictions. Dependence plots spotlighted metabolite interactions, such as the connection between hippuric acid and one of its derivatives in RCC, and between GM3(d34:1) and GM3(18:1_16:0) in OC, hinting at potential mechanistic relationships. Through decision plots, a detailed error analysis was conducted, contrasting feature importance for correctly versus incorrectly classified samples. In essence, our pipeline emphasizes the importance of harmonizing AutoML and XAI, facilitating both simplified ML application and improved interpretability in metabolomics data science.

**Availability:** https://github.com/obifarin/automl-xai-metabolomics

## Introduction

Machine learning (ML) has become a powerful tool in the field of non-targeted metabolomics, providing new ways to analyze and derive insight from data (Galal, et al., 2022; Liebal, et al., 2020). Metabolomics, the study of the small molecules produced by metabolism in biological systems, generates large amount of complex data (Ren, et al., 2015). This complexity emerges from the vast number of measured metabolites, the orders of magnitude of their concentrations, and the complex interconnected network of enzymatic reactions involved in metabolic pathways, necessitating advanced data analytics (Boccard and Rudaz, 2014). While there are many promising applications of ML in metabolomics, there are also challenges associated with its use, especially for non-experts. These mainly include the expertise required to select appropriate ML algorithms and to tune hyperparameters. In this respect, Automated ML (AutoML) methods have been proposed to improve the optimization workflow of creating ML models (Zöller and Huber, 2019).

AutoML tools enable automating the steps involved in ML (Zöller and Huber, 2019), reducing or even eliminating the need for expert knowledge in ML processes, which traditionally involve several specialized steps such as data preprocessing, feature engineering, model selection, and hyperparameter tuning. Numerous AutoML methods have been developed over time (He, et al., 2019). Tree based pipeline optimization (TPOT), for example, is an AutoML tool that uses genetic programming to automate the design of ML models by identifying the best data preprocessing steps, feature selection methods, and ML algorithms (Olson, et al., 2016). H2O AutoML abstracts away the complexities of developing the ML pipeline using supervised learning algorithms, ensemble learning (such as stacking and boosting), hyperparameter optimization (with random and grid search), and early stopping to improve predictive performance (LeDell and Poirier, 2020). Another popular AutoML framework is Auto-sklearn. This tool, based on the Scikit-learn library, automatically identifies and optimizes the appropriate ML algorithm and hyperparameters for a given dataset and task (Feurer, et al., 2015). It uses Bayesian optimization, meta-learning, and ensemble construction to create a powerful and robust automated ML solution (Feurer, et al., 2020; Feurer, et al., 2015).

Together with ML automation, the need for interpretable ML models and pipelines has also been increasing, especially in biomedicine (Vellido, 2020). This need has led to a rise in interest in explainable artificial intelligence (XAI) techniques, which seek to provide insights into how ML models make predictions (Rezaul Karim, et al., 2022). Two examples of this technique include Partial Dependence Plots (PDPs) and Individual Conditional Expectation (ICE) plots. PDPs show how the predicted outcome of a ML model changes as a given feature changes (Friedman, 2001). ICE plots, a variant of PDP, show how the predicted value changes as a feature value changes for each instance. This provides a more individualized interpretation than PDP, which shows the average change in predicted value for all instances (Goldstein, et al., 2013). These methods are global interpretation methods as they focus on explaining the behavior of the entire model. A technique that explains individual predictions (a local interpretation method) is Local Interpretable Model-agnostic Explanations (LIME) (Tulio Ribeiro, et al., 2016). LIME does this by locally approximating a complex model around the vicinity of a particular instance using a simpler model. This simpler model can then be used to explain the model’s prediction for that instance. Furthermore, Kernel SHAP (Shapley Additive Explanations) allows for both local and global interpretations (Lundberg and Lee, 2017). Kernel SHAP is a unified framework for interpreting model predictions across various ML algorithms, combining ideas from LIME and Shapley values to create a model-agnostic approach for approximating SHAP values. LIME explains “black-box” ML models by building interpretable local surrogate models, while Shapley values are a credit allocation scheme from cooperative game theory for feature importance attribution. Kernel SHAP has seen several applications across chemistry and biomedicine (Bloch, et al., 2021; Roder, et al., 2021; Rodriguez-Perez and Bajorath, 2020).

In this manuscript, we showcase a pipeline that combines both AutoML and XAI (AutoML-XAI) and apply it to two different metabolomics datasets. A seven urine metabolite discriminant panel for human renal cell carcinoma (RCC) (Bifarin, et al., 2021), and a seventeen-serum lipid discriminant panel for human ovarian cancer (OC) among Korean women (Sah, et al., 2023). Autosklearn is used to automate feature preprocessing, data preprocessing, and ML model selection, while Kernel SHAP is used as the XAI technique to study the model’s interpretability. Autosklearn was chosen partly because of its time hyperparameter that allows to control how long the AutoML will run, but also because of its ease of use with the popular ML library sklearn. Likewise, Kernel SHAP was chosen because of its utility as a model-agnostic XAI. Results demonstrate that Autosklearn outperforms standard ML models such as random forest, support vector machine (SVM), and k-nearest neighbors (kNN) for the metabolomics datasets under study. Moreover, Kernel SHAP allows for both global and local interpretations of the models, enabling the ranking of metabolomic feature importance. This can be visualized through summary and waterfall plots, respectively. Through dependence plots, XAI enables the assessment of how metabolomic features and their interactions with other features affect model predictions. Furthermore, decision plots enable a detailed error analysis of the AutoML model’s predictions.

## Methods

### Automated Machine Learning with Auto-Sklearn

The Auto-sklearn framework uses Bayesian optimization, meta-learning, and ensemble construction to automatically select ML pipelines (Feurer, et al., 2015). The Bayesian optimization step utilizes a probabilistic model to explore the space of possible hyperparameter settings, and to select the settings that are most likely to produce the best results, balancing ‘exploration’ with ‘exploitation’. The chosen hyperparameter setting is evaluated, the model is updated with the result, and the process is repeated. Meta-learning is used to warm-start the Bayesian optimization process. It allows Auto-sklearn to learn from previous experiences, and to use that knowledge to improve its performance on new tasks. In essence, Auto-sklearn uses meta-learning to reduce the search space and find optimal pipeline configurations more efficiently. The meta-features required for this task include information about the datasets, algorithms, and hyperparameters. Conversely, Auto-sklearn’s ensemble construction is based on ensemble selection, which entails generating many individual models and combining them to form an ensemble. The goal is to choose an ensemble that minimizes the cross-validation error on the training data.

### Local Interpretable Model-agnostic Explanations

LIME is an algorithm for explaining the predictions of any ML model (Tulio Ribeiro, et al., 2016). In brief, the algorithm works by building a simpler, interpretable model in the vicinity of the prediction of the black-box model. The interpretable model is then used to explain the prediction of the black-box model for an instance. The algorithm can be represented as follows:

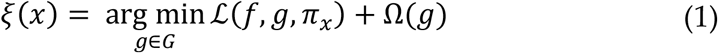

Where *x* is the instance to be explained, *g* is the interpretable model (e.g., linear regression), *G* is the family of interpretable models (e.g., linear models), *f* is the black box model to be explained, *π*_*x*_ is a proximity measure between a generated instance *z* and *x* (this is the kernel that defines locality), and *Ω*(*g*)is a regularization function. *L*(*f, g, π*_*x*_) is the locality-aware loss term that approximates *g* using *f*, as defined below:

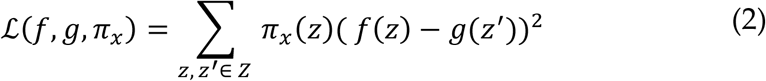

In essence, LIME leverages synthetic samples *Z*, generated by perturbing datasets *X*, to fit a local interpretable model. The objective is to train the interpretable model output *g*(*z*^′^) to match the output of the complex model *f*(*z*) (ground truth). Additionally, the samples are weighted by *π*_*x*_ to ensure the model is locally faithful. Hence, LIME constructs a linear model (or any other interpretable model) *g* in the vicinity of test instance, enabling meaningful local interpretations.

### Shapley Values

The Shapley value is a principled approach used to calculate the individual contributions of elements within a cooperative system (Lundberg and Lee, 2017). Shapley values are a fair way of determining the credit distribution for an outcome and have been widely used in game theory and cooperative game analysis.

Shapley values are formally represented as:

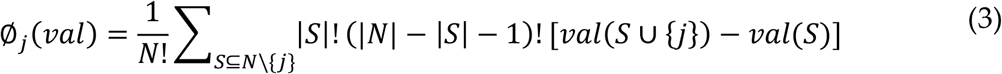

Where ∅_*j*_(*val*) is the Shapley value of element *j, N* is the number of elements in the cooperative system, *S* is the subset of elements in the system and *val*(*S* ∪ {*j*}) − *val*(*S*) represents the marginal contribution of element *j*. Shapley values are desirable for feature attribution in ML as they consider the marginal contribution of each feature to the final output, rather than considering the contribution of a feature in isolation, providing a more comprehensive attribution of the contribution of each feature to the outcome.

Shapley values satisfy several desirable axioms, such as additivity, symmetry, and null effect property, making them a robust and flexible method for feature attribution. The additivity axiom states that the sum of the Shapley values of the individual elements in a cooperative system is equal to the total value of the system. For ML applications, this guarantees local feature attribution as well as global interpretations. Additivity is formally expressed as:

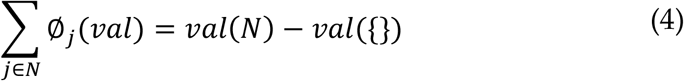

On the other hand, the symmetry axiom refers to the property that the value assigned to an element by the Shapley value is the same, no matter the order in which the element is added to the coalition. In ML, Shapley values can be used to fairly distribute the contribution of each feature to the prediction made by a model. For two elements *j* and *k*, the symmetry axiom can be represented as:

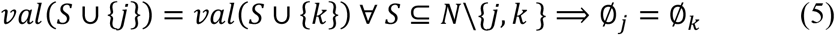

Lastly, the dummy axiom guarantees that if an element does not have any impact on a cooperative system, the Shapley value will be null. For an element *z*, the dummy axiom can be represented as:

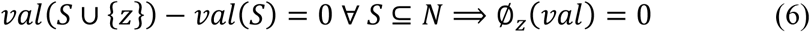

Shapley values provide a robust and fair way of attributing the contribution of each feature to the outcome, their mathematical properties making them an ideal tool for feature attribution in ML.

### Kernel SHAP

Kernel SHAP is a method for estimating Shapley values for any ML model (Lundberg and Lee, 2017). It does this by using a weighted linear regression, where the weights are defined by a specially-designed kernel function. The goal is to explain the output of the function *f*(*x*) as the sum of the contributions of each feature value in *x* (additivity property).

The feature attribution function *f*(*S*) is typically defined as the expected output of the model given that the feature values in *S* are known and the other features are ‘missing’. However, directly evaluating this function for all subsets of features is generally not feasible because of the combinatorial explosion of possible subsets and the need to retrain the model on each subset.

To address this issue, Kernel SHAP uses a LIME-inspired approach that selects a small, representative sample of feature subsets. These are then used to estimate a simpler, more interpretable function *g* that approximates the complex model in the neighborhood of a specific prediction. The approximation is constructed using a weighted linear regression, where the weights are determined by a kernel function *π*_*x*_:

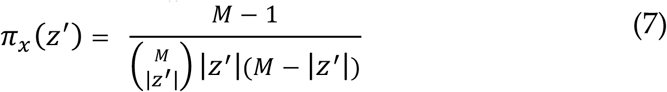

Where *z*^′^ is a binary vector indicating the presence or absence of features in the simplified instance *x*^′^. The maximum coalition size is represented by *M*, while |*z*^′^| is the number of present features in *z*^′^. The solution to the optimization problem for constructing *g* to approximate the true model *f* in the vicinity of the instance *x*, is as follows:

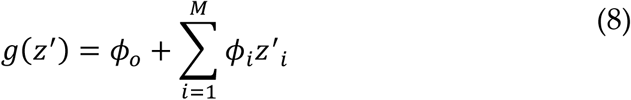

The linear model *g* is trained by optimizing the following loss function ℒ:

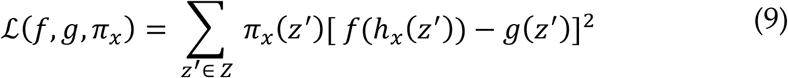

Where the training data is represented by *z*^′^, *h*_*x*_(.) maps the coalition vector to feature values, *f*(.) is the black box model, and *g*(.) is the linear model.

### Machine Learning Pipeline

One of the datasets utilized for this work was derived from a urine-based metabolomics study that proposed a metabolite panel that discriminates between renal cell carcinoma (RCC) patients (*n* = 82) and healthy controls (*n* = 174), with a focus on specific urine biomarkers (*n*=7) (Bifarin, et al., 2021). The second dataset is derived from the serum lipidomic analysis of ovarian cancer (OC) patients of Korean descent, encompassing diverse histological types and disease stages (*n* = 208), compared with women with other gynecological malignancies, including invasive cervical cancer (*n* =117) (Sah, et al., 2023). For this work, we focused on a 17-lipid panel that distinguished OC from non-OC patients.

Python’s Pandas library (version 2.0.0) was used for data handling and the numpy library (version 1.23.5) was used for numerical computations. For the RCC study, to facilitate the subsequent model training and testing, the data were randomly partitioned into a training set (80% of the data) and a testing set (20% of the data). In the case of OC study, the data split followed the strategy used in the published work: the dataset was divided into a training set with 70% (OC *n* = 144 and non-OC *n* = 83) and a test set with 30% (OC *n* = 64 and non-OC *n* = 34) of the samples.

The training set was subsequently balanced (OC *n* = 144 and non-OC *n* = 144) using the Synthetic Minority Over-sampling Technique (SMOTE). All datasets were autoscaled after the partitioning. For the classification problem, ML models from Scikit-learn library (version 0.24.2) were used with default parameters, including Random Forest Classifier (RF), Support Vector Classifier (SVC), and k-Nearest Neighbors Classifier (*k*-NN), alongside the AutoML approach *via* autosklearn (version 0.15.0). Model performance was assessed using the Receiver Operating Characteristic Area Under the Curve (ROC AUC), accuracy, sensitivity, and specificity score.

For AutoML, AutoSklearnClassifier was allowed a total of 600 seconds to identify the optimal ML pipelines in the case of the RCC dataset, and 3600 seconds in the case of OC dataset. The resampling strategy used was cross-validation with default parameters using the hold out strategy, which splits the training data further into an internal training and validation set using a 67:33 ratio. The ensemble method was used, and up to 50 models were considered for inclusion in the ensemble. The amount of time that each model was allowed to run during the optimization process was 240 seconds for the RCC dataset, and 1440 seconds for the OC dataset. Following model training, the data was extracted from the trained AutoSklearnClassifier using PipelineProfiler (version 0.1.18), allowing for the visualization of the pipeline matrix.

Finally, for comparing model performance after autosklearn built the AutoML pipelines, the final ensemble was fitted to the entire training dataset under 5-fold cross validation and tested on the unseen datasets, in addition to RF, SVC, and *k*-NN. In the second part of the ML analysis, SHAP (version 0.41.0) was used for XAI. A summary of the training dataset was created using *k*-means clustering (*k*=10) to feed into the SHAP KernelExplainer, which computed SHAP values for each feature and each instance in the test set. The SHAP values were then used to create summary, beeswarm, waterfall, force, dependence, and decision plots for detailed explanations. An overview of this approach is presented in **Figure 1**, while a synopsis of the ML pipeline is given in **Figure 2**.

**Figure 1.**
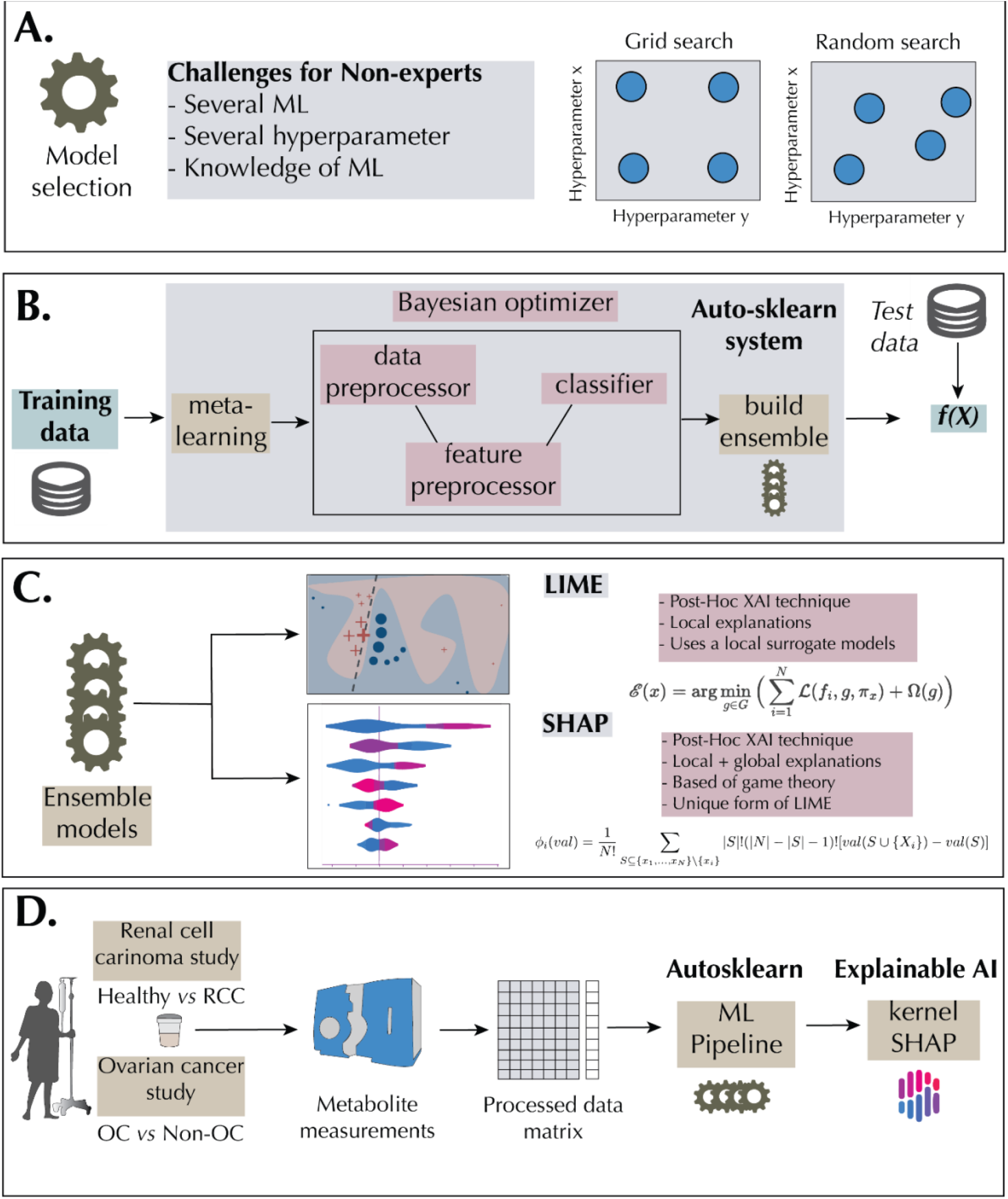
Automated ML-Explainable AI Workflow. (A) Highlight of the challenges associated with ML model selection for non-experts. Grid and random searches are typically leveraged for hyperparameter selection. (B) The Auto-Sklearn AutoML system is based on meta-learning, Bayesian optimization, and ensemble construction. (C) Ensemble models constructed *via* Auto-Sklearn can be interpreted with Explainable AI (XAI) techniques such as LIME and SHAP. (D) Application of AutoML and XAI to a RCC urine and OC serum metabolomics dataset. Local Interpretable Model-agnostic Explanations – LIME, Shapley Additive exPlanation – SHAP, Renal Cell Carcinoma – RCC, Ovarian Cancer– OC.

**Figure 2.**
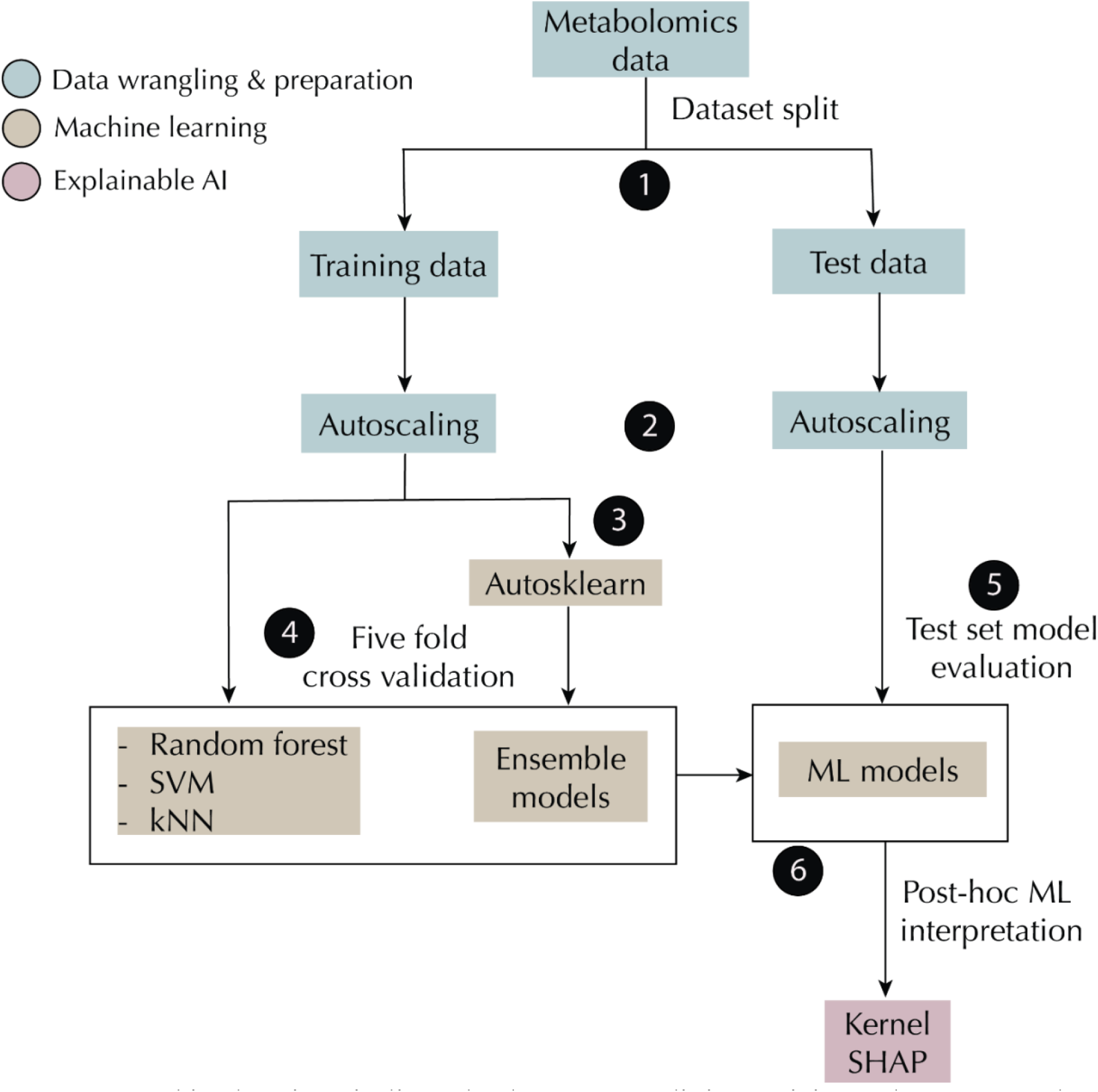
Machine learning pipeline. The dataset was split into training and test sets, each was subsequently auto-scaled. ML models were built using the training set and their performances were accessed using the test set. AutoML ensemble models were explained using kernel SHAP.

## Results

### Model Evaluation and Performance

Using the metabolomics datasets, RF, SVM, and k-NN algorithms were compared to AutoML *via* autosklearn on a test dataset following model training. For the RCC dataset, Auto-sklearn yielded the best results, achieving an AUC score of 0.97 (training score: 0.97 ± 0.03), an accuracy score of 0.90 (training score: 0.96 ± 0.02), and a sensitivity score of 0.89 (training score: 0.88 ± 0.06). The RF model matched Auto-sklearn in accuracy, registering a test score of 0.90 and a training score of 0.95 ± 0.03. Meanwhile, both the SVM and RF models surpassed Auto-sklearn only in specificity, each recording a test set score of 0.94 (SVM training score: 0.99± 0.01, RF training score: 0.99± 0.03). Similarly, for the OC dataset, Auto-sklearn yielded the best results, achieving an AUC score of 0.85 (training score: 0.86 ± 0.07), an accuracy score of 0.78 (training score: 0.79 ± 0.09), a sensitivity score of 0.75 (training score: 0.74 ± 0.06), and a specificity score of 0.82 (training score: 0.82 ± 0.09). Furthermore, the RF model matched Auto-sklearn in only AUC (training score: 0.86± 0.07, test score: 0.85) and sensitivity (training score: 0.81± 0.02, test score: 0.75). In brief, on most of the metrics considered, autosklearn had superior classification performance. For the complete machine learning performance results, see **Table 1**.

**Table 1:** Machine learning performance for RCC and OC metabolomics datasets. The training set results are indicated by the score (±standard deviation). The best test set score for each metric is in bold and underlined. Metrics reported include Receiver Operating Characteristic Area Under the Curve (ROC AUC), accuracy, sensitivity, and specificity. *k*-NN: K-nearest neighbors. SVM: Support Vector Machines.

| Model | Train set AUC | Test set AUC | Train set Accuracy | Test set Accuracy | Train set Sensitivity | Test set Sensitivity | Train set Specificity | Test set Specificity |
| --- | --- | --- | --- | --- | --- | --- | --- | --- |
| **Renal Cell Carcinoma** |  |  |  |  |  |  |  |  |
| Random Forest | 0.97 (0.03) | 0.95 | 0.95 (0.03) | **<u>0.90</u>** | 0.88 (0.06) | 0.83 | 0.99 (0.03) | **<u>0.94</u>** |
| SVM | 0.97 (0.03) | 0.93 | 0.92 (0.03) | 0.90 | 0.77 (0.08) | 0.83 | 0.99 (0.01) | **<u>0.94</u>** |
| *k*-NN | 0.96 (0.04) | 0.93 | 0.94 (0.04) | 0.87 | 0.83 (0.09) | 0.78 | 0.99 (0.01) | 0.91 |
| AutoML | 0.97 (0.03) | **<u>0.97</u>** | 0.96 (0.02) | **<u>0.90</u>** | 0.88 (0.06) | **<u>0.89</u>** | 0.99 (0.02) | 0.91 |
| **Ovarian Cancer** |  |  |  |  |  |  |  |  |
| Random Forest | 0.86 (0.07) | **<u>0.85</u>** | 0.80 (0.04) | 0.74 | 0.81 (0.02) | **<u>0.75</u>** | 0.79 (0.09) | 0.74 |
| SVM | 0.84 (0.04) | 0.82 | 0.76 (0.02) | 0.73 | 0.78 (0.05) | 0.72 | 0.75 (0.06) | 0.76 |
| *k*-NN | 0.77 (0.05) | 0.69 | 0.69 (0.07) | 0.63 | 0.57 (0.07) | 0.56 | 0.82 (0.06) | 0.76 |
| AutoML | 0.87 (0.06) | **<u>0.85</u>** | 0.79 (0.09) | **<u>0.78</u>** | 0.74 (0.06) | **<u>0.75</u>** | 0.82 (0.09) | **<u>0.82</u>** |

### AutoML Selected Configurations

The AutoML approach for the RCC dataset had 289 algorithms run, with only four of these runs failing due to crashes or exceeding the time limit. For the OC dataset, autosklearn had 1373 algorithms run with only 22 runs failing for similar reasons. **Figure 3A & B** show the pipeline profile for RCC and OC datasets, respectively. In the RCC dataset, the primitive contribution shows that the Extra trees classifier and Random trees embeddings, which generates embeddings using decision trees, were associated with high ROC AUC scores, while kernel PCA was associated with low scores. Similarly, in the OC dataset (**Fig 3B**), gradient boosting and feature agglomeration were associated with high ROC AUC scores, while, as in the RCC pipeline, kernel PCA was associated with low scores. **Figure 3C** shows a typical pipeline graph, starting with the input dataset, followed by class balancing with weighting where the under-represented class is given a higher weight while the over-represented class is given a lower weight. This makes the classifier focus on the minority class during training. After balancing, random trees embeddings was used as a feature preprocessor. Finally, the embeddings are passed onto a SVM for the classification task.

**Figure 3.**
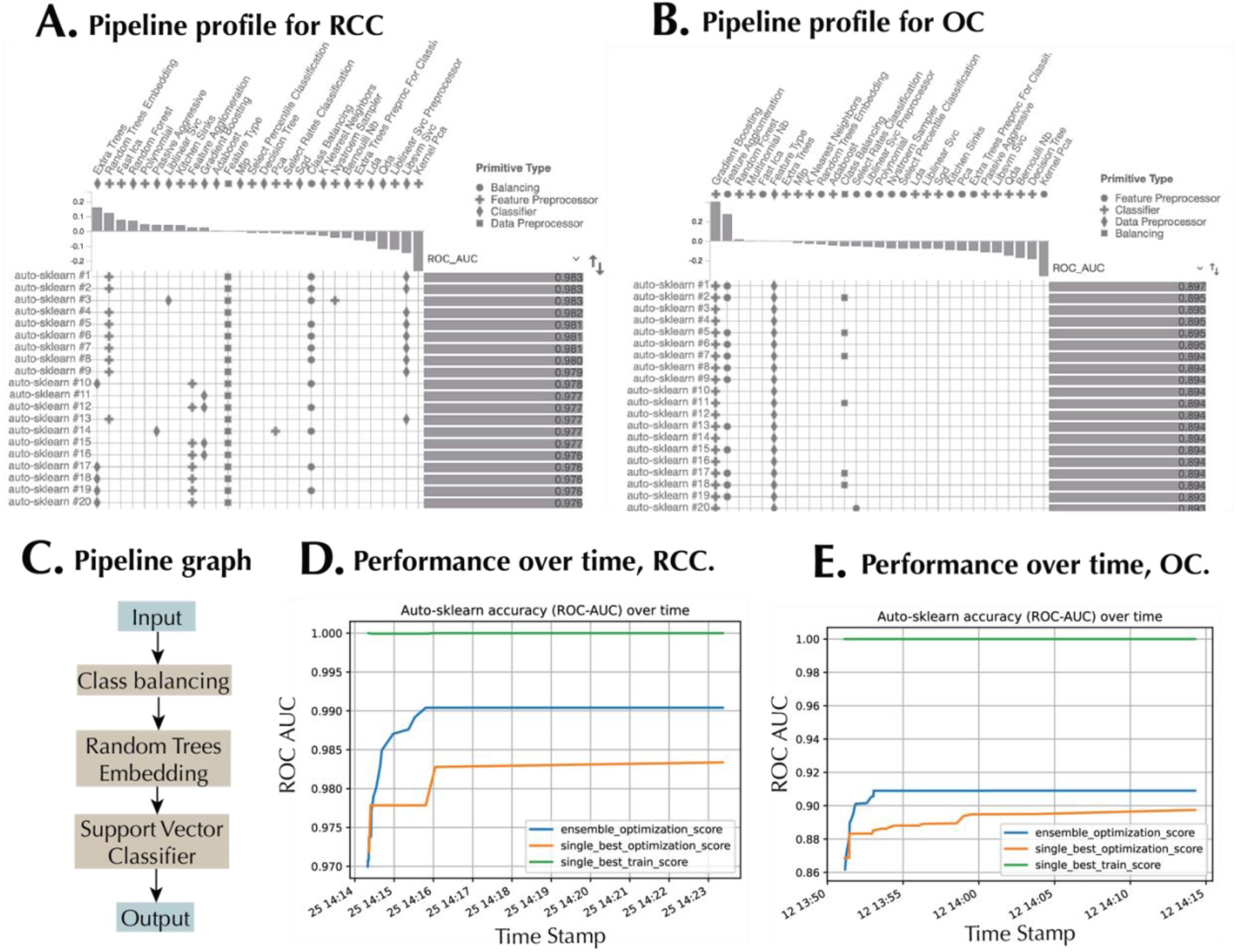
Automated machine learning pipelines. Pipeline profile showing the pipeline primitives, pipeline matrix, and the corresponding ROC AUC scores for the (A) RCC dataset, and (B) OC dataset. Only 20 successful ML pipelines are shown in each case. (C) Pipeline graph for a sample autoML pipeline. ML pipeline performance over time during model training for the (D) RCC dataset, and (E) OC dataset. The scores reported include the single best score on the internal training set, the single best optimization score, and the ensemble optimization score.

**Figure 3D & E** shows the performance of autosklearn over time during the RCC and OC dataset training sessions, respectively. Three scores were used. The ensemble optimization score reflects the performance of the ensemble of models on the internal validation data. The single best optimization score represents the performance of the top-performing individual model on the validation data, while the single best train score shows the fitting of the best single model on the training set (a subset of the entire training set). Higher ensemble optimization score compared to the single best optimization score revealed the benefit of using an ensemble selection strategy.

The final ensemble model for the RCC dataset consisted of 13 ML pipelines with the following classifiers selected: SVM with a linear kernel (liblinear_svc), SVM with a non-linear kernel option (libsvm_svc), gradient boosting, and ExtraTrees classifier, a meta estimator that fits several randomized decision trees (extra_trees). The feature preprocessors selected in the configurations included Nystroem Sampler (approximates kernel maps for SVMs), random trees embeddings (generates embeddings using decision trees), and feature agglomeration (clusters features for dimensionality reduction). For the balancing strategy, weighing was used (**Table 2**). Similarly, the OC ensemble model comprised 16 ML pipelines, all of which employed gradient boosting as their chosen classifier. The selected feature pre-processor includes feature agglomeration and Select Rates Classification. As for the balancing strategy, it also involved the utilization of weighting (**Table 2**).

**Table 2:**
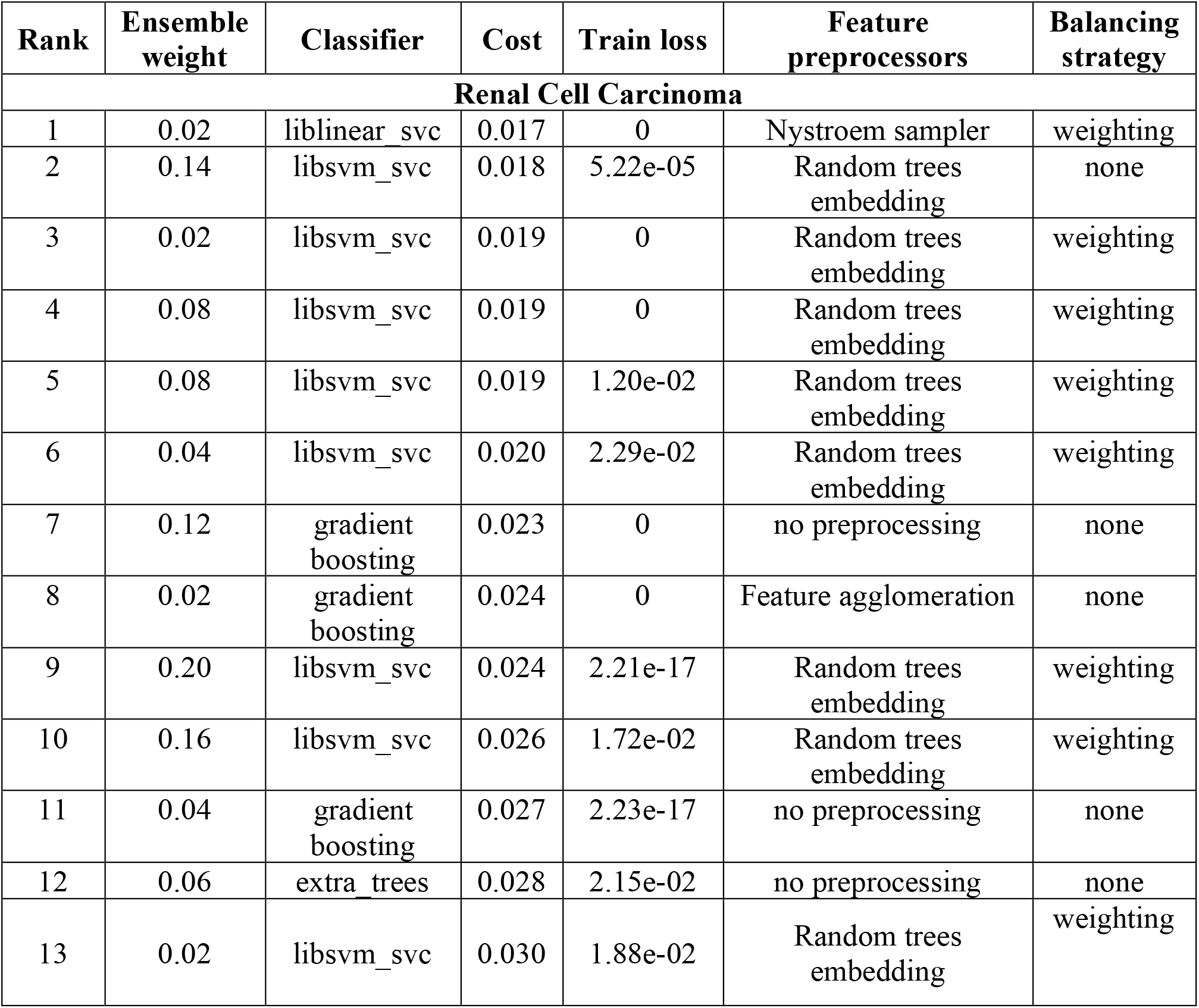
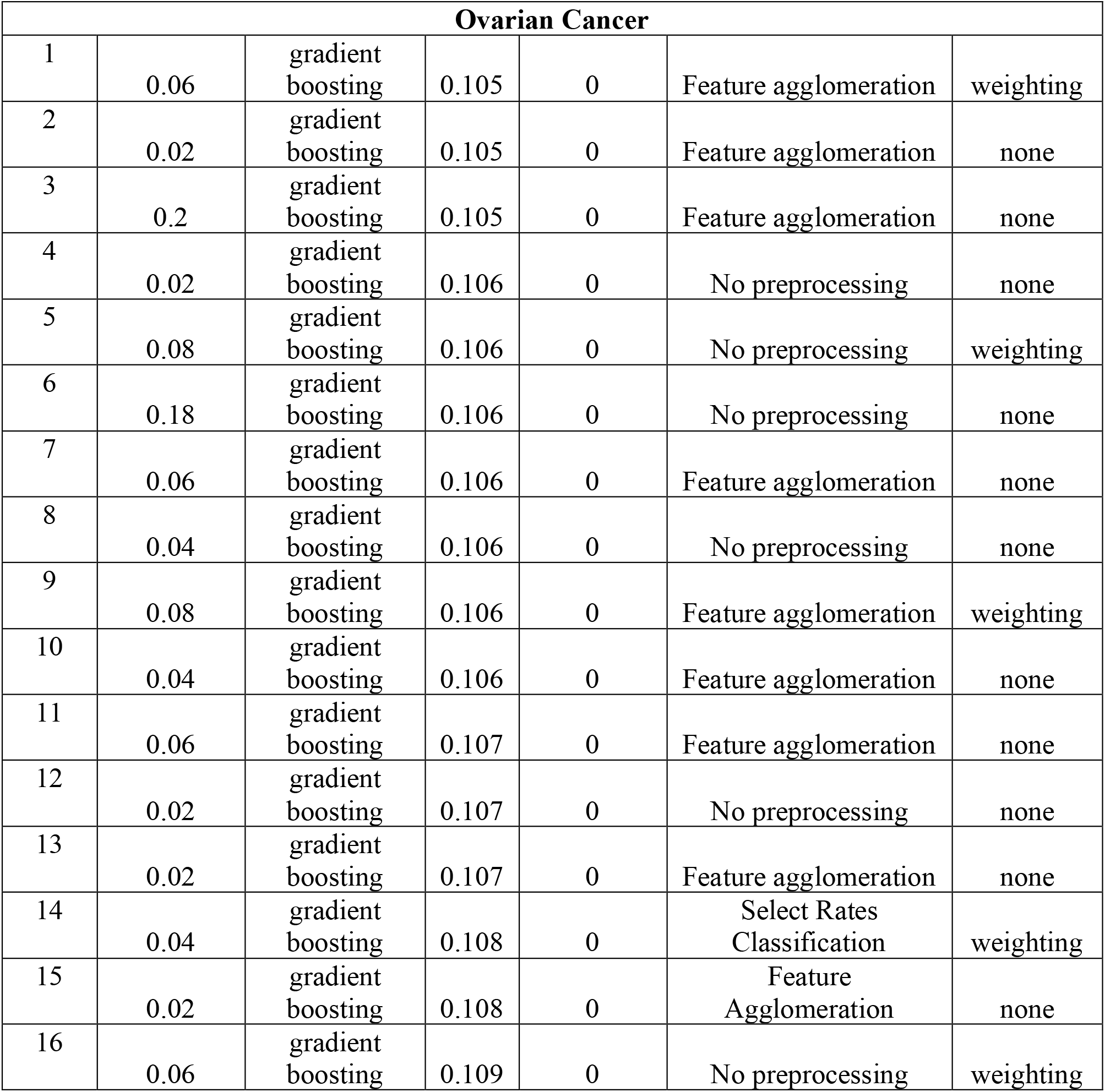
Final ensemble model used for the RCC and OC dataset. The rank of the model is based on its “cost”. The cost is the loss of the model on the validation set, while the train loss is the loss of the model on the training set. Weighting gives higher weight to the under-represented classes during training. The ensemble weight is the weight of the ML pipeline in the final ensemble.

For the RCC dataset, Random trees embedding and Libsvm SVC were associated with higher ensemble weight, while Liblinear SVC and feature agglomeration were associated with lower ensemble weight for the final ensemble generated (**Figure S1A**). On the other hand, Nystroem Sampler and Liblinear SVC were associated with high ROC AUC while Extra Trees were associated with low ROC scores within the ensemble model (**Figure S1B**). Furthermore, in the case of the OC dataset, class balancing was associated with higher ensemble weights, while select rates classification was associated with lower ensemble weights (**Figure S1C**). In the case of ROC AUC for the OC dataset, feature agglomeration and select rates classification are associated with higher and lower values, respectively (**Figure S1D**). Overall, the AutoML method had the best performance on the test set, leveraging the diversity of ML primitives and ensemble construction.

### Interpretation of AutoML Pipeline Ensemble *via* SHAP

SHAP values measure the impact of each metabolite on the disease status prediction for a given sample, providing a measure of each feature importance. **Figure 4A** shows the beeswarm or summary plot for the RCC dataset, with each point representing a Shapley value *ϕ*_*i,j*_ where *i* are the samples and *j* are the metabolites. The color of each point indicates the relative abundance of a metabolite, with darker colors representing higher values. For example, dibutylamine has the highest range of Shapley values, making it the most important feature for discriminating between RCC and healthy controls. Low dibutylamine urine levels are associated with lower Shapley values, leading to a more likely patient classification as a healthy control. On the other hand, N-acetyl-glucosaminic acid is the least important feature for discriminating between RCC and healthy controls in this panel, with small Shapley values in both positive and negative directions.

**Figure 4.**
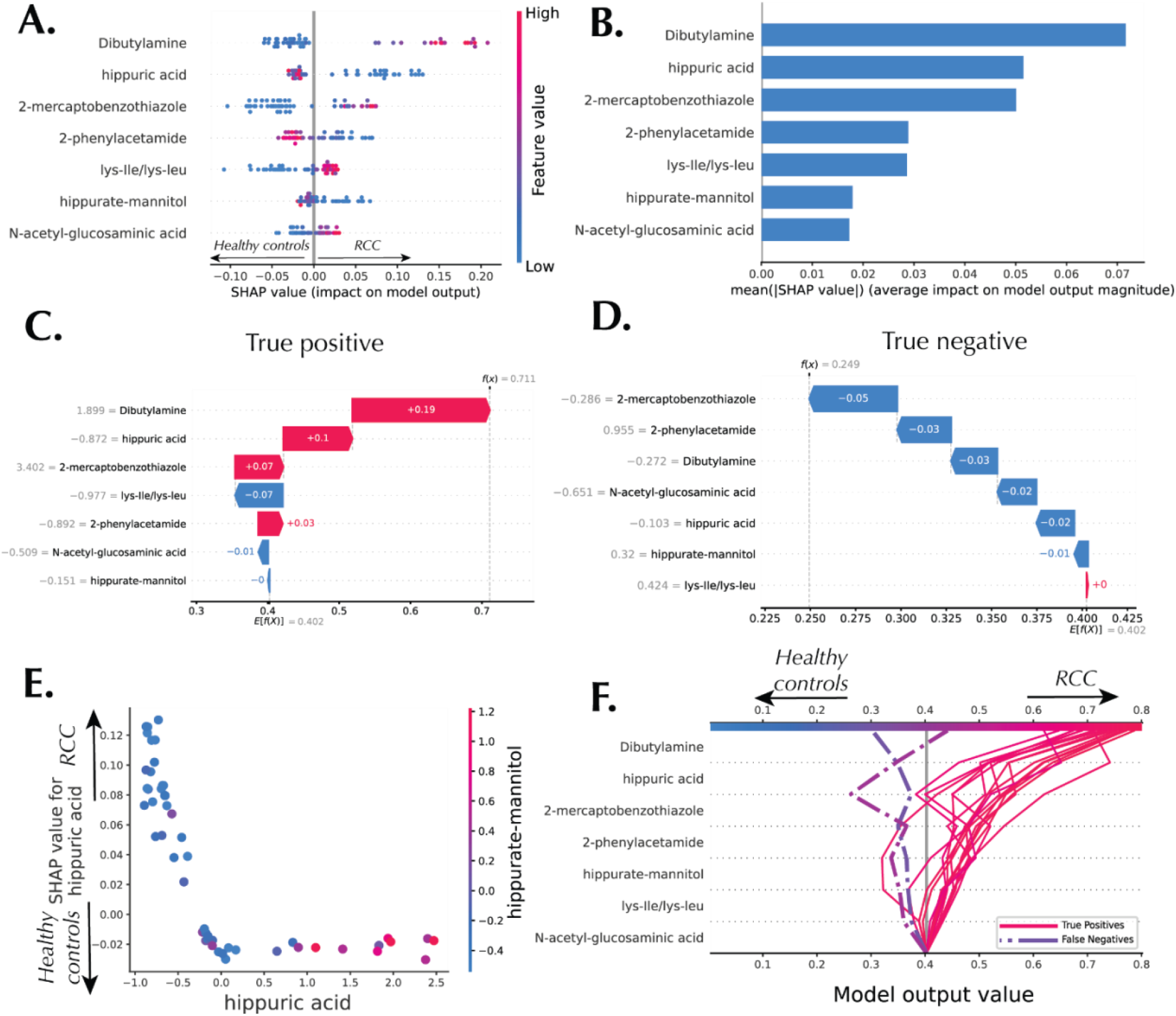
Machine learning interpretations of the ensemble model constructed by AutoML for the RCC dataset. (A) Beeswarm plot and (B) Summary plot showing global interpretation of the model. (C) Waterfall plot – local explanation for a true positive (RCC) sample. (D) Waterfall plot **–** local explanation for a true negative (healthy control) sample. (E) Dependence plot showing the interaction between hippuric acid and the hippurate-mannitol derivative. (F), Decision plot highlighting true positive and false negative samples.

**Figure 4B** shows the importance plot, a less detailed version of beeswarm plot focused on the feature importance ranking for each of the metabolites in the RCC dataset. **Figure 4C & D** show the waterfall plot for representative true positive and true negative urine samples. Shapley values *ϕ*_*i,j*_ are represented as arrows that either increase or decrease the prediction *f*(*x*_*i*_) when compared to the expected prediction E*f*(*x*). For example, for the true positive sample presented in **Figure 4C**, the predicted RCC probability value is 0.71, with dibutylamine, hippuric acid, 2-mercaptobenzothiazole, and 2-phenylacetamide having a positive effect on the RCC prediction outcome, while Lys-Ile/Lys-Leu had the largest negative effects on prediction outcome. On the other hand, the true negative sample shown in **Figure 4D** had virtually all metabolites contributing towards a healthy prediction outcome, leading to an output of 0.25.

The SHAP dependence plot (**Figure 4E**) shows how the Shapley values for hippuric acid change with its relative abundance, showing a somewhat L-shaped pattern hinting at the presence of an inflection point occurring around the mean of 0 in how the metabolite abundance affects the model output. This points to the model’s fundamental dependence on whether the abundance of hippuric acid is low or high relative to the data distribution. Furthermore, the color of the plot is based on the relative abundance of another metabolite – tentatively identified as hippurate-mannitol, a derivative of hippuric acid. The behavior of this metabolite maps on to the patterns of hippuric acid as shown in the dependence plot. In **Figure 4F** the decision plots for all samples predicted as RCC is shown. This SHAP plot illustrates one or multiple predictions by visualizing the SHAP feature contributions for each of the samples. They demonstrate how the model combines feature evidence to make the RCC decision. The plot shows the features ranked in increasing order of importance starting from the bottom of the graph.

Similar interpretations were computed for the OC dataset. The summary plot (**Figure 5A**) shows that GM3(d34:1) is the most important metabolite for discriminating between non-OC and OC patients, followed by LPC(14:0), GM1(18:1_16:0), GM3(18:0_24:2), *etc*. The local interpretation of a true positive sample (a correctly classified OC patient) and a true negative sample (a correctly classified non-OC patient) are shown in **Figures 5B & C**, respectively. The dependence plot in **Figure 5D** shows the interaction between two ganglioside GM3 family species: GM3(d34:1) and GM3(18:1_16:0). High abundance of both lipids drives OC classification, and vice versa. The decision plot of all samples classified as OC is presented in **Figure 5E**.

**Figure 5.**
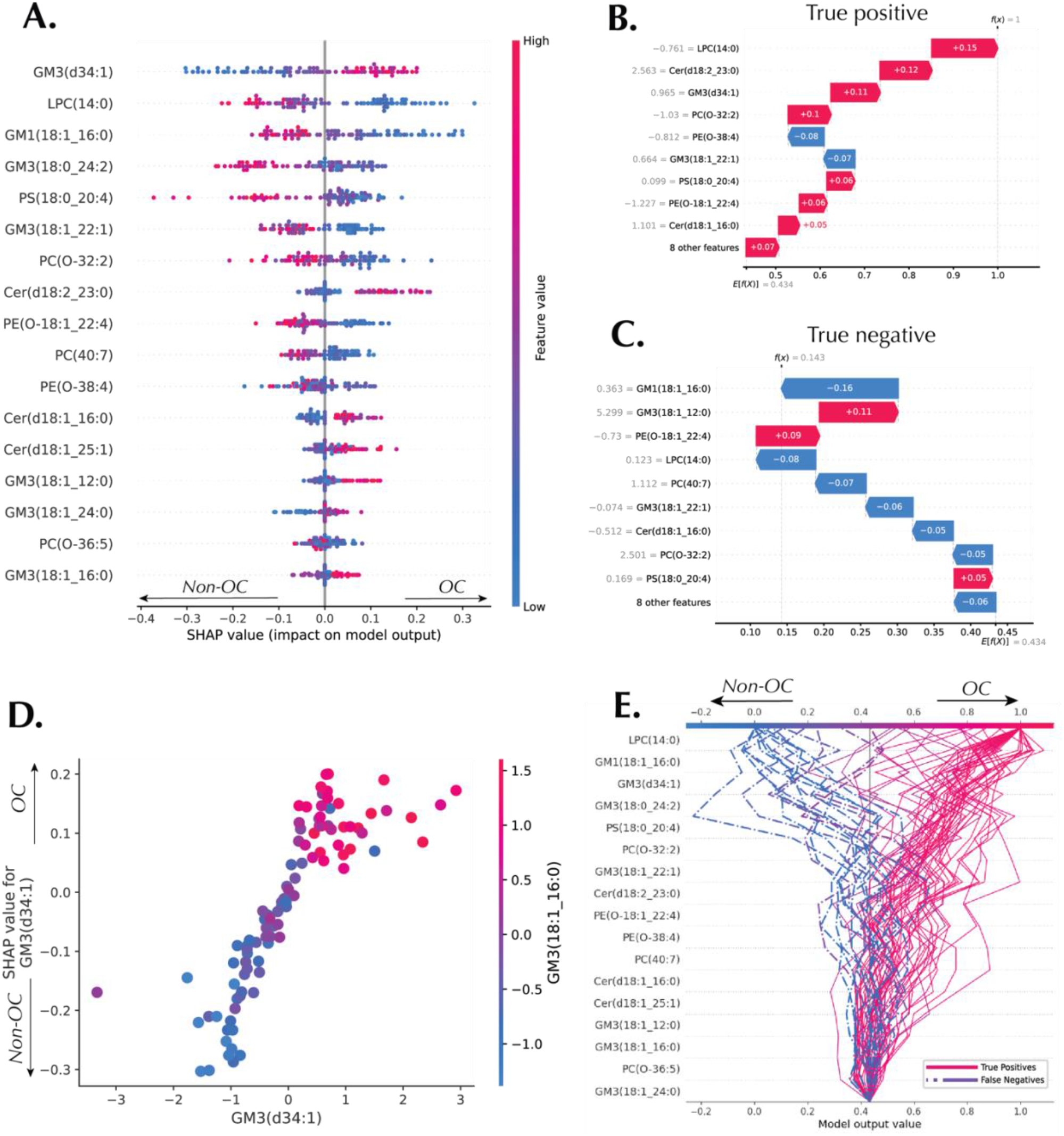
Machine learning interpretations of the ensemble model constructed by AutoML for the OC dataset. (A) Beeswarm plot and (B) Waterfall plot – local explanation for a true positive (OC) sample. (C) Waterfall plot **–** local explanation for a true negative (non-OC) sample. (D) Dependence plot showing the interaction between GM3(d34:1) and the GM3(18:1_16:1). (E), Decision plot highlighting true positive and false negative samples for OC and non-OC classification.

The RCC decision plots for true negative samples (**Figure 6A**) and false positive samples (**Figure 6B**) have, expectedly, different orders of feature ranking. For detailed error analysis powered by SHAP, the feature importance rankings between true negatives (TN) and false positives (FP) were compared. Features were ranked by their average SHAP values in each group (**Figures 6A&B**), and Kendall’s Tau correlation coefficient between the two rankings were computed to quantify the consistency of feature importance across groups (**Figure 6C**, *τ* = −0.14, *p* = 0.7). **Figure 6D** shows the change in feature important rank between true negative and false positive samples, with Lys-Ile/Lys-Leu having the highest displacement in ranking from the second most important metabolite in the TN samples to the least important feature in FP samples. RCC decision plots for all true positive (TP) and false negative (FN) samples are presented in **Figures 6E&F**. The comparison of feature importance consistency in all predicted RCC samples (TP *vs*. FN) is presented in **Figure 6G** (*τ* = 0.62, *p* = 0.07). These results indicate a better consistency between the predicted RCC samples (TP *vs*. FN) in comparison with the predicted healthy controls (TN *vs*. FP). N-acetyl glucosaminic acid had the highest displacement in ranking from the least important metabolite in TP samples to the fourth most important metabolite in FN samples. Other metabolites, except for dibutylamine, had only one place displacements in feature importance ranking (**Figure 6H**). Similar error analysis was conducted for the OC diagnostic model, with results presented in **Figures S2** and S**3**.

**Figure 6.**
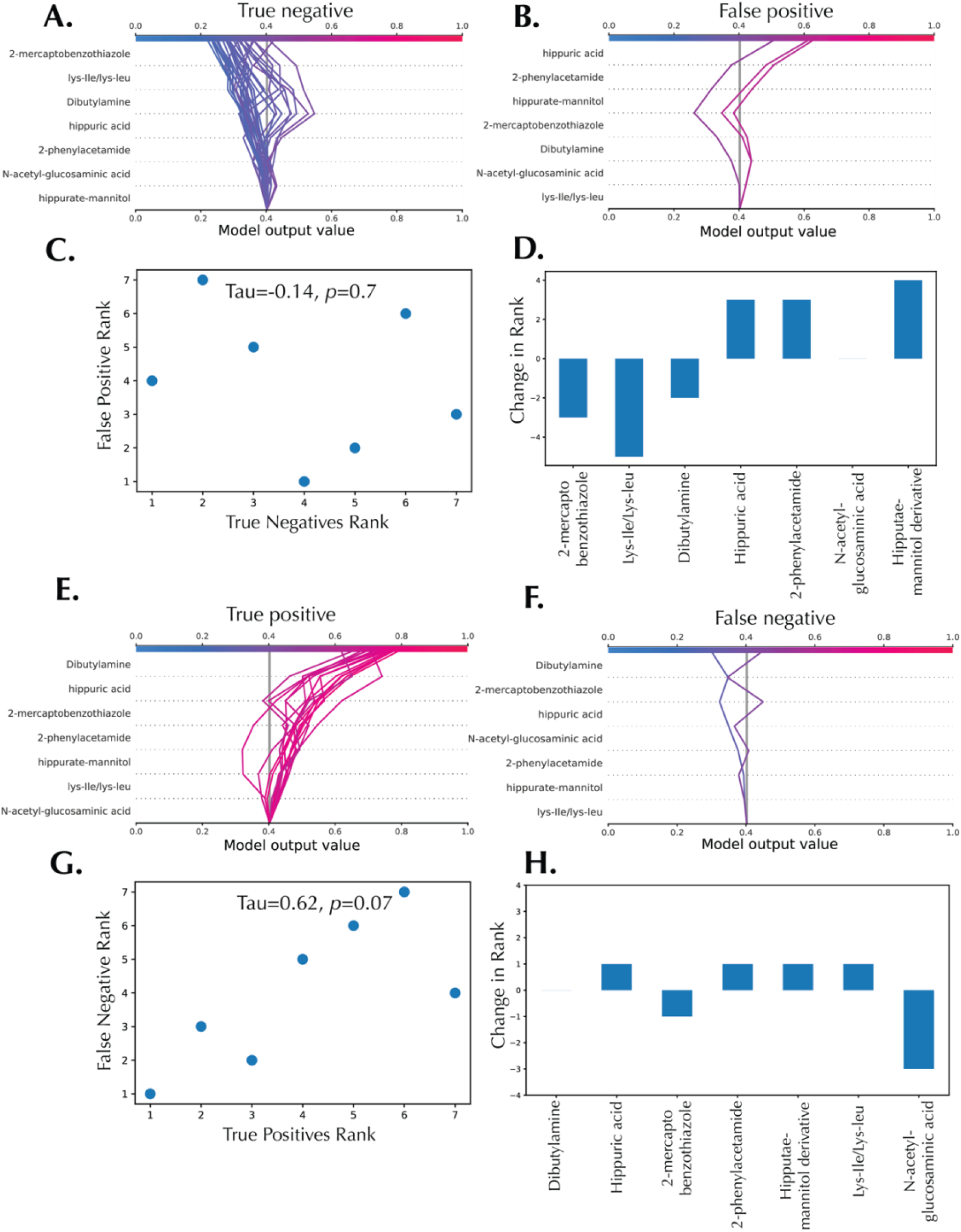
Error analysis decision plots for the RCC diagnostic model. (A) Decision plot for all true negative samples. (B) Decision plot for all false positive samples. (C) Feature importance rank correlation between true negative and false positive samples. (D) Changes in feature importance rank between true negatives *vs*. false positives. (E) Decision plot for all true positive samples. (F) Decision plot for all false negative samples. (G) Feature importance rank correlation between true positive and false negative samples. (H) Changes in feature importance rank between true positive *vs*. false negative. *Tau* is Kendall’s Tau correlation coefficient.

## Discussion

In this study, we demonstrated a comprehensive computational pipeline that effectively combines AutoML and XAI for the analysis of cancer metabolomics data. We focused our application on a urine-based metabolomics dataset with 7 biomarkers, aimed at distinguishing RCC patients from healthy controls (Bifarin, et al., 2021) and 17 serum lipid markers for discriminating between OC and non-OC patients (Sah, et al., 2023). The use of ML in metabolomics has been promising for biomarker discovery and understanding of biological systems (Mendez, et al., 2019), but its complexity, especially for non-experts, has been a barrier to widespread adoption. The use of AutoML simplifies the ML process (Feurer, et al., 2020; Feurer, et al., 2015). Furthermore, the addition of an XAI technique to the pipeline provides more interpretable results, making it a valuable approach for building hypothesis based on the discovery data.

We utilized Auto-sklearn, a robust and flexible AutoML tool, to automate the ML process (Feurer, et al., 2015). Auto-sklearn uses Bayesian optimization, meta-learning, and ensemble construction to automatically select the best ML pipelines, including data preprocessing, feature engineering, model selection, and hyperparameter tuning. We compared the performance of Autosklearn with traditional ML models such as Random Forest, SVM, and k-NN. The AutoML ensemble, leveraging diverse algorithms like SVM, gradient boosting, and Extra Trees classifier, outperformed standalone models, achieving the best ROC AUC of 0.97 on the RCC test set. In the case of the OC, gradient boosting was selected as the classifier for all pipelines. However, different feature preprocessors were utilized for various pipelines. This resulted in optimum performance scores for AUC, accuracy, sensitivity, and specificity. These results not only highlight the efficacy of AutoML methods such as Auto-sklearn in simplifying intricate ML tasks, but also underscore the superior performance achievable that stems from the use of ensemble construction.

To interpret the created AutoML model, we used Kernel SHAP, an XAI technique (Lundberg and Lee, 2017). Kernel SHAP offers both global and local interpretations of the model, allowing for an understanding of each metabolite contribution to the prediction. Shapley values facilitated feature importance ranking, revealing dibutylamine as the most discriminative feature for RCC detection. Meanwhile, GM3(d34:1) emerged as the most significant metabolite for differentiating non-OC from OC patients. In addition to global interpretations, waterfall plots provided local explanations on metabolite effects on individual predictions. Dependence plots in the RCC study showed interactions between metabolites like hippuric acid and hippurate-mannitol, a derivative, possibly indicating interconnected biological pathways or processes. In the case of the OC dataset, the interaction between GM3(d34:1) and GM3(18:1_16:1) was highlighted. Such insights uncover mechanistic relationships and new hypotheses meriting further investigation.

Detailed error analysis was conducted to compare feature rankings between correctly and incorrectly classified samples. The results were presented with decision plots, which provide a clear description of which metabolites had a significant shift in their importance between these two groups, showing how the model’s reliance on certain metabolites varies depending on whether a given sample is correctly or incorrectly classified. Interestingly, in the RCC study, metabolome feature rank consistency was higher between true positives and false negative (*τ*=0.62, *p*=0.07) *vs*. true negative and false positive (*τ*=-0.14, *p*=0. 7). This might be partly explained by the fact that there are less false negatives (n=2) compared to false positives (n=3).

In conclusion, our pipeline exemplifies a successful integration of automated ML and model interpretability for metabolomics datasets, reducing the complexities of advanced ML application for non-experts and leveraging XAI for bolstering model transparency. By providing an evaluation of various ML models and an intricate explanation of the auto-sklearn model’s decision-making process, our work not only enhances the interpretability and trustworthiness of predictive diagnostic models but also highlights the transformative potential of AutoML and XAI in computational metabolomics.

## Acknowledgements

FMF acknowledges support by NIH 1R01CA218664-01.

## Supplementary Information

**Figure S1.**
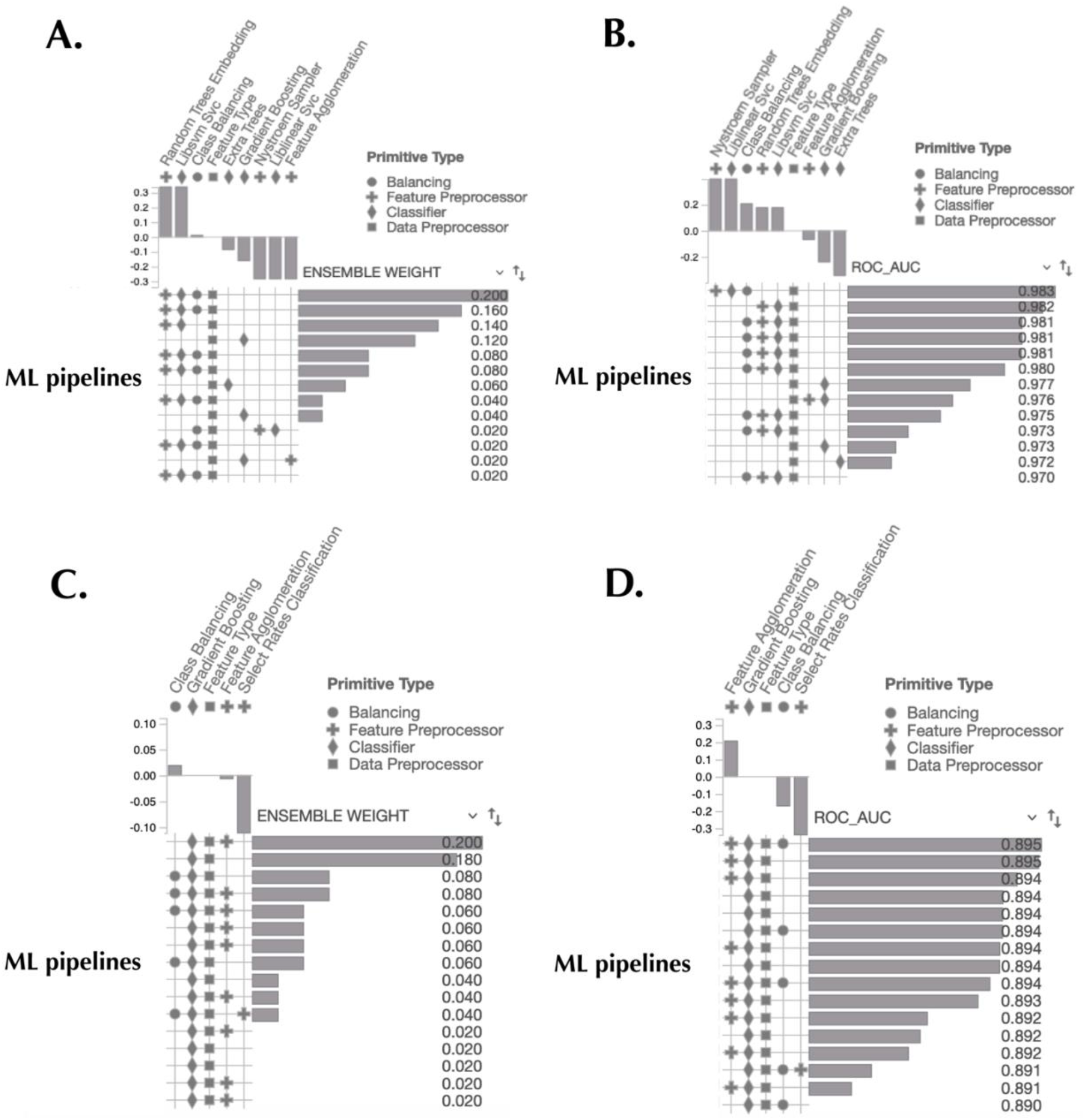
Pipeline profile for the ensemble model. (A) ML pipelines showing the ensemble weight contributions, RCC dataset. (B) ML pipelines showing the ROC AUC score ranking for each pipeline, RCC dataset. (A) ML pipelines showing the ensemble weight contributions, OC dataset. (B) ML pipelines showing the ROC AUC score ranking for each pipeline, OC dataset.

**Figure S2.**
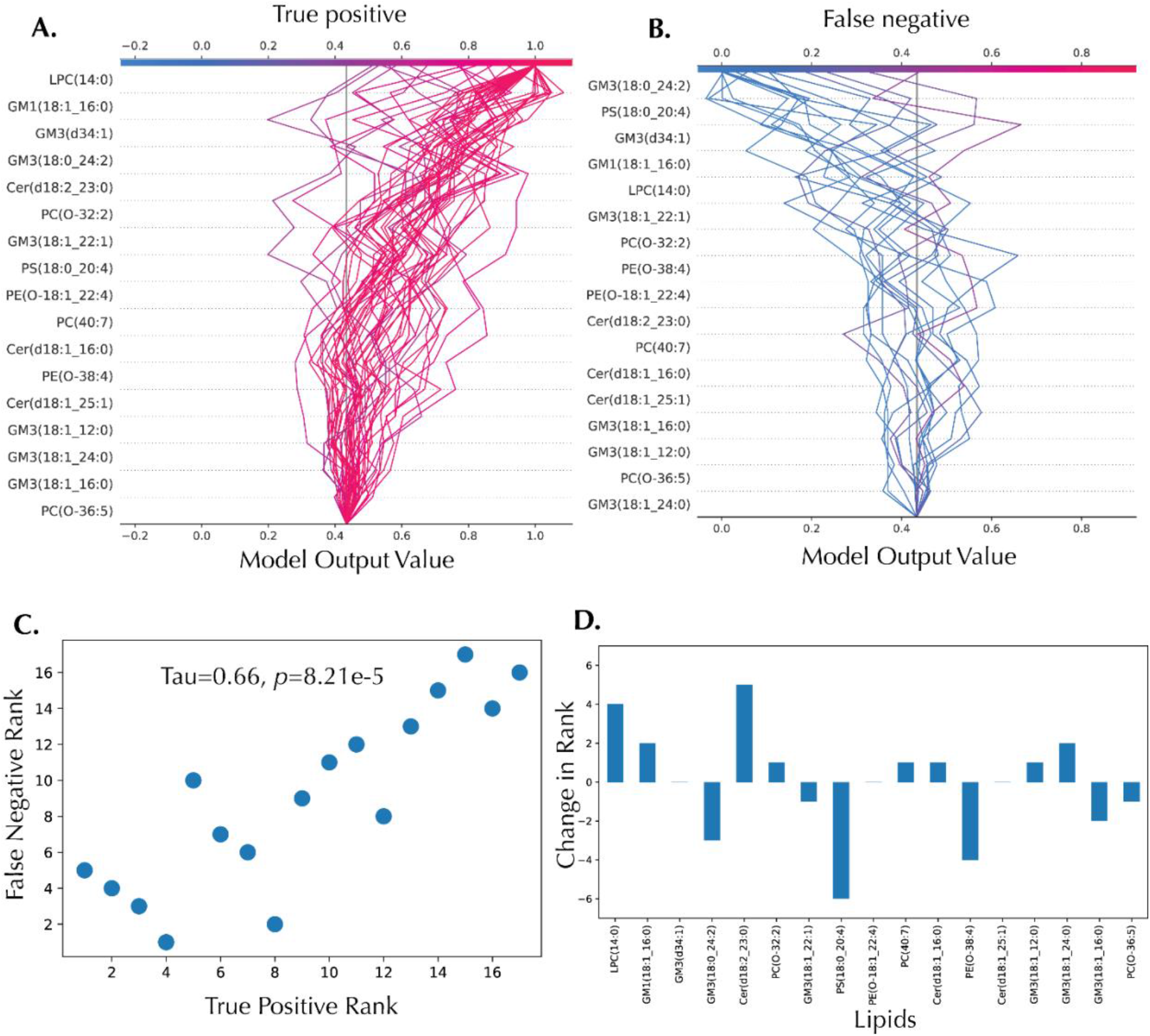
Error analysis decision plots for the OC diagnostic model, true positives *vs*. false negatives. (A) Decision plot for all true positive samples. (B) Decision plot for all false negative samples. (C) Feature importance rank correlation (D) Changes in feature importance rank.

**Figure S3.**
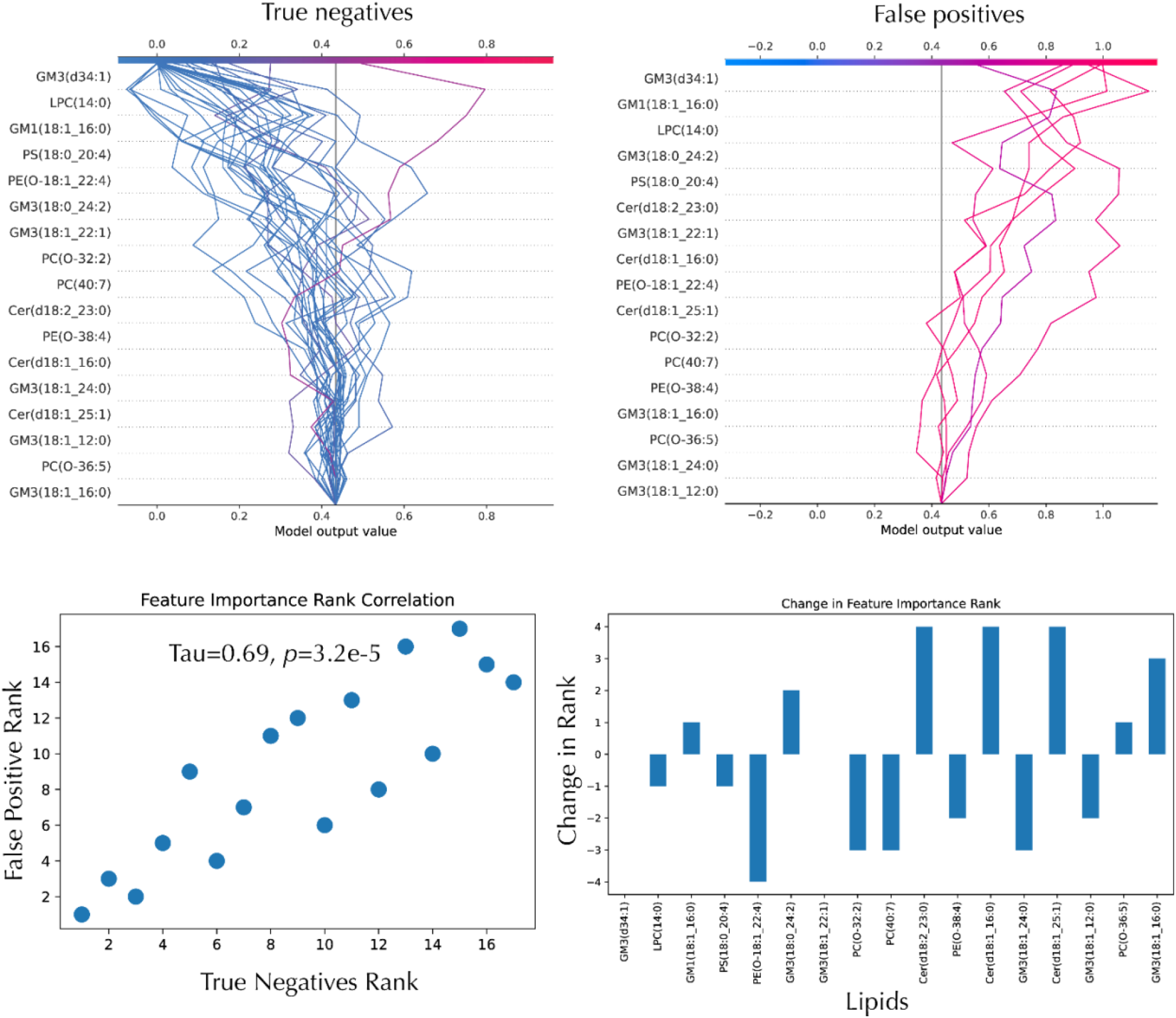
Error analysis decision plots for the OC diagnostic model, true negatives *vs*. false positives. (A) Decision plot for all true negative samples. (B) Decision plot for all false positive samples. (C) Feature importance rank correlation (D) Changes in feature importance rank.

## References

Bifarin, O.O., et al. Machine Learning-Enabled Renal Cell Carcinoma Status Prediction Using Multiplatform Urine-Based Metabolomics. J Proteome Res 2021;20(7):3629–3641.

Bloch, L., Friedrich, C.M. and Alzheimer’s Disease Neuroimaging, I. Data analysis with Shapley values for automatic subject selection in Alzheimer’s disease data sets using interpretable machine learning. Alzheimers Res Ther 2021;13(1):155.

Boccard, J. and Rudaz, S. Harnessing the complexity of metabolomic data with chemometrics. Journal of Chemometrics 2014;28(1):v–v.

Feurer, M., et al. Auto-Sklearn 2.0: Hands-free AutoML via Meta-Learning. In.; 2020. p. arXiv:2007.04074.

Feurer, M., et al. Efficient and Robust Automated Machine Learning. Neural Information Processing Systems 28 2015.

Friedman, J.H. Greedy Function Approximation: A Gradient Boosting Machine The Annals of Statistics 2001;29(5):1189–1232.

Galal, A., Talal, M. and Moustafa, A. Applications of machine learning in metabolomics: Disease modeling and classification. Front Genet 2022;13:1017340.

Goldstein, A., et al. Peeking Inside the Black Box: Visualizing Statistical Learning with Plots of Individual Conditional Expectation. In.; 2013. p. arXiv:1309.6392.

He, X., Zhao, K. and Chu, X. AutoML: A Survey of the State-of-the-Art. In.; 2019. p. arXiv:1908.00709.

LeDell, E. and Poirier, S. H2O AutoML: Scalable Automatic Machine Learning. 7th ICML Workshop on Automated Machine Learning 2020.

Liebal, U.W., et al. Machine Learning Applications for Mass Spectrometry-Based Metabolomics. Metabolites 2020;10(6).

Lundberg, S. and Lee, S.-I. A Unified Approach to Interpreting Model Predictions. In.; 2017. p. arXiv:1705.07874.

Mendez, K.M., Reinke, S.N. and Broadhurst, D.I. A comparative evaluation of the generalised predictive ability of eight machine learning algorithms across ten clinical metabolomics data sets for binary classification. Metabolomics 2019;15(12):150.

Olson, R.S., et al. Evaluation of a Tree-based Pipeline Optimization Tool for Automating Data Science. In.; 2016. p. arXiv:1603.06212.

Ren, S., et al. Computational and statistical analysis of metabolomics data. Metabolomics 2015;11:1492–1513.

Rezaul Karim, M., et al. Explainable AI for Bioinformatics: Methods, Tools, and Applications. In.; 2022. p. arXiv:2212.13261.

Roder, J., et al. Explaining multivariate molecular diagnostic tests via Shapley values. BMC Med Inform Decis Mak 2021;21(1):211.

Rodriguez-Perez, R. and Bajorath, J. Interpretation of machine learning models using shapley values: application to compound potency and multi-target activity predictions. J Comput Aided Mol Des 2020;34(10):1013–1026.

Sah, S., et al. Serum Lipidome Profiling Reveals a Distinct Signature of Ovarian Cancer in Korean Women. In.: bioRxiv; 2023.

Tulio Ribeiro, M., Singh, S. and Guestrin, C. “Why Should I Trust You?”: Explaining the Predictions of Any Classifier. In.; 2016. p. arXiv:1602.04938.

Vellido, A. The importance of interpretability and visualization in machine learning for applications in medicine and health care. Neural Computing and Applications 2020;32:18069– 18083.

Zöller, M.-A. and Huber, M.F. Benchmark and Survey of Automated Machine Learning Frameworks. In.; 2019. p. arXiv:1904.12054.

